# Metastatic potentials classified with hypoxia-inducible factor 1 downstream gene in pan-cancer cell lines

**DOI:** 10.1101/2023.05.08.539810

**Authors:** Kazuya Nakamichi, Kentaro Semba, Jun Nakayama

## Abstract

Hypoxia-inducible factor 1 (HIF1) gene codes a transcription factor that is stabilized under hypoxia conditions via post-translational modifications. HIF1 regulates tumor malignancy and metastasis by gene transcriptions, such as Warburg effect- and angiogenesis-related genes, in cancer cells. However, the HIF1 downstream genes show varied expressional patterns in different cancer types. Herein, we performed the hierarchical clustering based on the HIF1 downstream gene expression patterns using 1,406 cancer cell lines crossing 30 types of cancer to understand the relationship between HIF1 downstream genes and the metastatic potential of cancer cell lines. Four types of cancer were classified by HIF1 downstream genes with significantly altered metastatic potentials. Furthermore, different HIF1 downstream gene subsets were extracted to discriminate each subtype for the four cancer types. HIF1 downstream subtyping classification will help understand the novel insight into tumor malignancy and metastasis in each cancer type.

**Funding:** This work was supported by Project for JSPS KAKENHI (Grant-in-Aid for Scientific Research (C): JP23K06665 to JN, Grant-in-Aid for Early-Carrier Scientists: JP21K15562 to JN), and in part by translational research program from Fukushima Prefecture (KS).

**Competing Interests statement:** The authors have declared that no conflict of interest exists.

## Introduction

Hypoxia-inducible factor 1 (HIF1) is a heterodimeric transcription factor composed of HIF1α and HIF1β. HIF1α proline residues are hydroxylated by prolyl hydroxylase domain-containing protein under normoxia conditions [1], causing HIF1 degradation via ubiquitination by the E3 ubiquitin ligase containing von Hippel-Lindau protein [2]. Conversely, the HIF1 protein becomes stable and regulates the expression of target genes under hypoxia conditions.

HIF1 contributes to tumor malignancy by regulating cancer metabolism, microenvironment, and tumor stemness by regulating downstream genes [3–5], including vascular endothelial growth factor, aldehyde dehydrogenase, carbonic anhydrase 9, and glucose transporter 1 [6–9]. Especially, HIF1 is involved in multiple steps of metastasis, including epithelial-mesenchymal transition (EMT), invasion, angiogenesis, intravasation, extravasation, and pre-metastatic niche formation. For example, HIF1 directly regulates EMT-related gene expression, such as Zinc Finger E-Box Binding Homeobox 1 [10], Snail Family Transcriptional Repressor 1 [11], and Twist Family BHLH Transcription Factor 1 [12], invasion-related gene expressions, such as Lysyl Oxidase, Lysyl Oxidase Like 2 [13], and angiogenesis gene expression, such as VEGFA [14]. Conversely, HIF1 downstream genes are heterogeneous at the expressional level with tumor microenvironmental stress [15].

Therefore, we focused on the diversity of HIF1 downstream genes and cancer metastasis. First, a hierarchical clustering analysis of cancer cell lines which were available in the public dataset, cancer cell line encyclopedia (CCLE), was performed to reveal HIF1 downstream gene diversity. Four cancer types were identified and classified using HIF1 downstream genes. Next, the metastatic potential scores of cancer cell lines were subjected to metastatic organ tropism and metastatic activity analyses. The classified groups of four cancer types revealed the differences in metastatic potential among the classified groups. These findings support the importance of understanding the HIF1 downstream in terms of metastasis and prognosis.

## Results

### Flowchart of the classification with a public database

We obtained the gene expression data [16] and metastatic potential score data [17] for cancer cell lines by CCLE. A total of 1,406 cell lines from CCLE are divided into 30 lineages. The gene expression data of 1,406 cell lines were analyzed by hierarchical clustering in each lineage. Cell lines were classified into two clusters in 24 lineages. The other 6 lineages were not divided into clusters. The metastatic potential score was then used to evaluate the two clusters in 24 lineages. A two-tailed Mann–Whitney U test was used to reveal the significant difference in 4 out of 24 lineages between the two groups. Finally, we extracted differential expressed genes whose |log fold-change (FC) | was over 1 (Figure 1).

**Figure 1.**
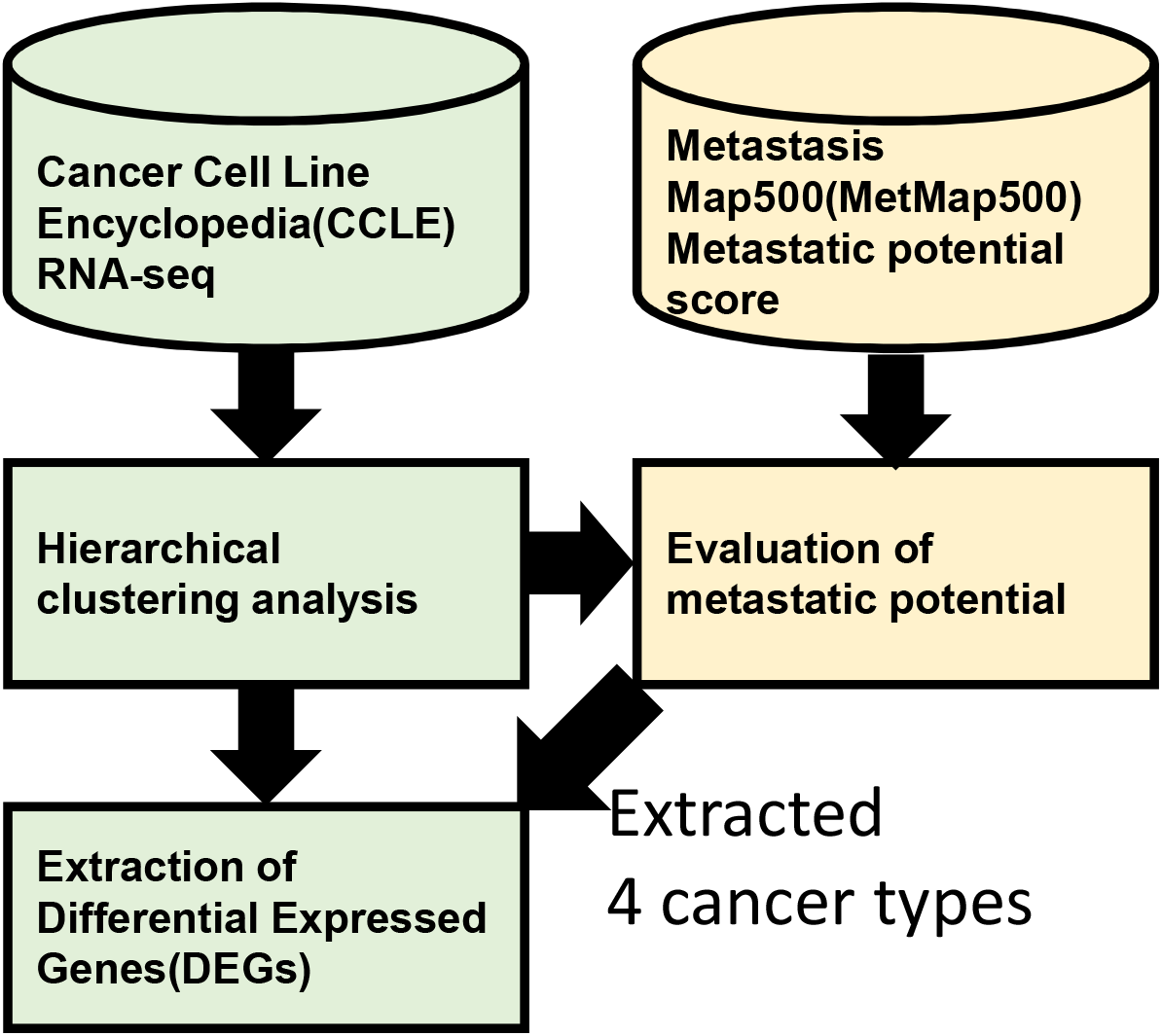
Flowchart of clustering with HIF1 downstream genes. The gene expression data of the 1,406 cell lines were analyzed by hierarchical clustering in each lineage. Cell lines were classified into two clusters in 24 lineages. Then, the two clusters in 24 lineages were evaluated by metastatic potential score. A significant difference was found in 4 out of 24 lineages between the two groups using a two-tailed Mann–Whitney U test. Finally, we extracted DEGs whose |log FC| was over 1.

### Classification by HIF1 downstream genes with a metastatic score

Hierarchical clustering with metastatic potential by HIF1 downstream genes was performed as shown in the flowchart of Figure 1. First, we selected the gene list [18] from MSigDB [19] as HIF1 downstream genes in this analysis. Second, RNA-seq data of HIF1 downstream genes in the 1,406 cell lines were analyzed by hierarchical clustering in each lineage. Of the total 30 lineages, 24 were classified into two major groups. The two divided groups were categorized as “Group A” and “Group B.”.” Group A in bone tumor contained 16 cell lines of osteosarcoma and 3 cell lines of chondrosarcoma, while Group B contained only Ewing sarcoma, considering the cell lines in detail (Figure S1). Additionally, this result suggests differential HIF1 downstream gene expression between Ewing sarcoma and osteosarcoma/chondrosarcoma. Group A contained 17, 13, and 6 cell lines of Luminal-like, human epidermal growth factor receptor 2 positive, and Basal-like subtypes in breast cancer, respectively. Conversely, Group B contained cell lines with only Basal-like subtypes. Uterine cancer classification highlighted a subtype with Group B that is mainly composed of endometrial adenocarcinoma (Figure S1). Then, a two-tailed Mann–Whitney U test was used to evaluate the metastatic potential score [17] between Group A and Group B. The result revealed significant differences in metastatic potential scores in 4 lineages out of 24 lineages. In bone tumors, 19 and 21 cell lines were to Groups A and B, respectively (Figure 2A and Table S1). Group B had significantly higher metastatic potential to the liver than Group A (p = 0.02214) (Figure 3A and Table S2). In breast cancer, 37 and 26 cell lines belonged to s A and B, respectively (Figure 2B and Table S1). Group B had significantly higher metastatic potential to the bone and liver than Group A (*p* = 0.01778 and 0.01471) (Figure 3B, and Table S2). In upper aerodigestive cancer, 22 and 34 cell lines belonged to Groups A and B, respectively (Figure 2C and Table S1). Group B had significantly higher metastatic potential to the liver than Group A (*p* = 0.02461) (Figure 3C and Table S2). In uterine cancer, 13 and 27 cell lines belonged to Group A and B, respectively (Figure 2D and Table S1). Group B had significantly higher overall metastatic potential than Group A (*p* = 0.02496) (Figure 3D and Table S2). Other cancer and metastatic destinations did not make significant differences between the two groups.

**Figure 2.**
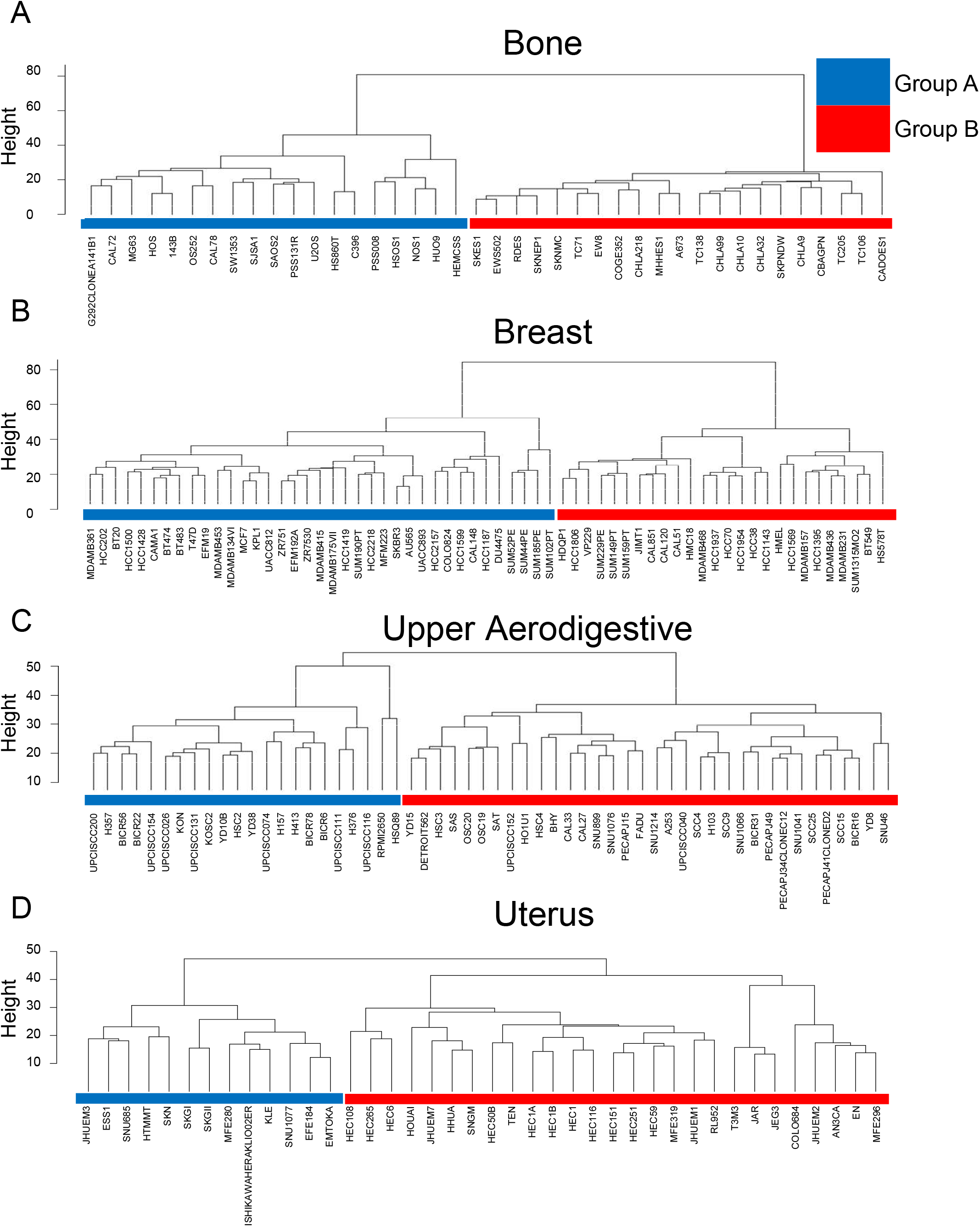
Reclassification of four types of cancer cell lines with hierarchical clustering using HIF1 downstream genes. Hierarchical clustering classified cancer cell lines into two groups. Group A was shown in blue label while Group B in red label.

**Figure 3.**
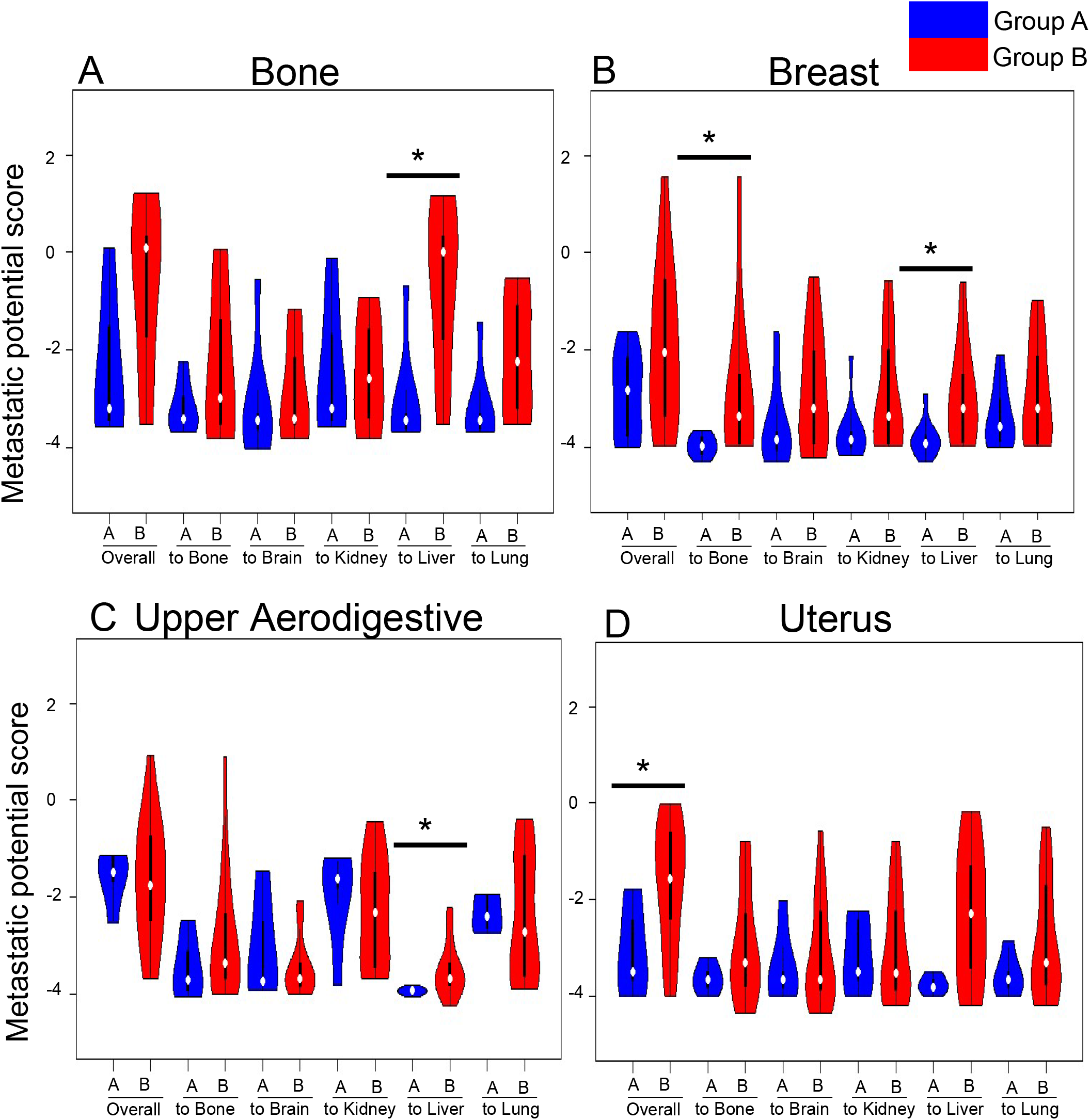
Metastatic potentials in reclassified cancer cell lines. Metastatic potential scores for each metastatic organ in Groups A and B were shown by Violin plots, with Group A shown in blue and Group B in red. \**P* < 0.05

Next, we extracted differential expressed genes (DEGs) to obtain genes effective for classification. We calculated the logFC of each gene between Groups A and B. The genes with |logFC|>1 were regarded as DEGs. In bone tumors, 36 genes were more highly expressed in Group A, while 14 genes were more highly expressed in Group B (Figure 4A, S2A, and Table S3). In breast cancer, 9 genes were more highly expressed in Group A, while 26 genes were more highly expressed in Group B (Figure 4B, S2B, and Table S3). In upper aerodigestive cancer, none of the genes were more highly expressed in Group A, while 19 genes were more highly expressed in Group B (Figure 4C, S2C, and Table S3). In uterine cancer, 20 genes were more highly expressed in Group A, while 3 genes were more highly expressed in Group B (Figure 4D, S2D, and Table S3). Venn diagrams were drawn for each of the upregulated genes in Groups A and B to study overlap from the comparative analysis of DEGs in high metastatic group (Figure 4E, 4F, and Table 1). In all four cancer types, none of the common genes were upregulated or downregulated in the highly metastatic group. Only caveolin 1 (CAV1) increased in the high metastatic group in three cancer types (Figure 4F).

**Figure 4.**
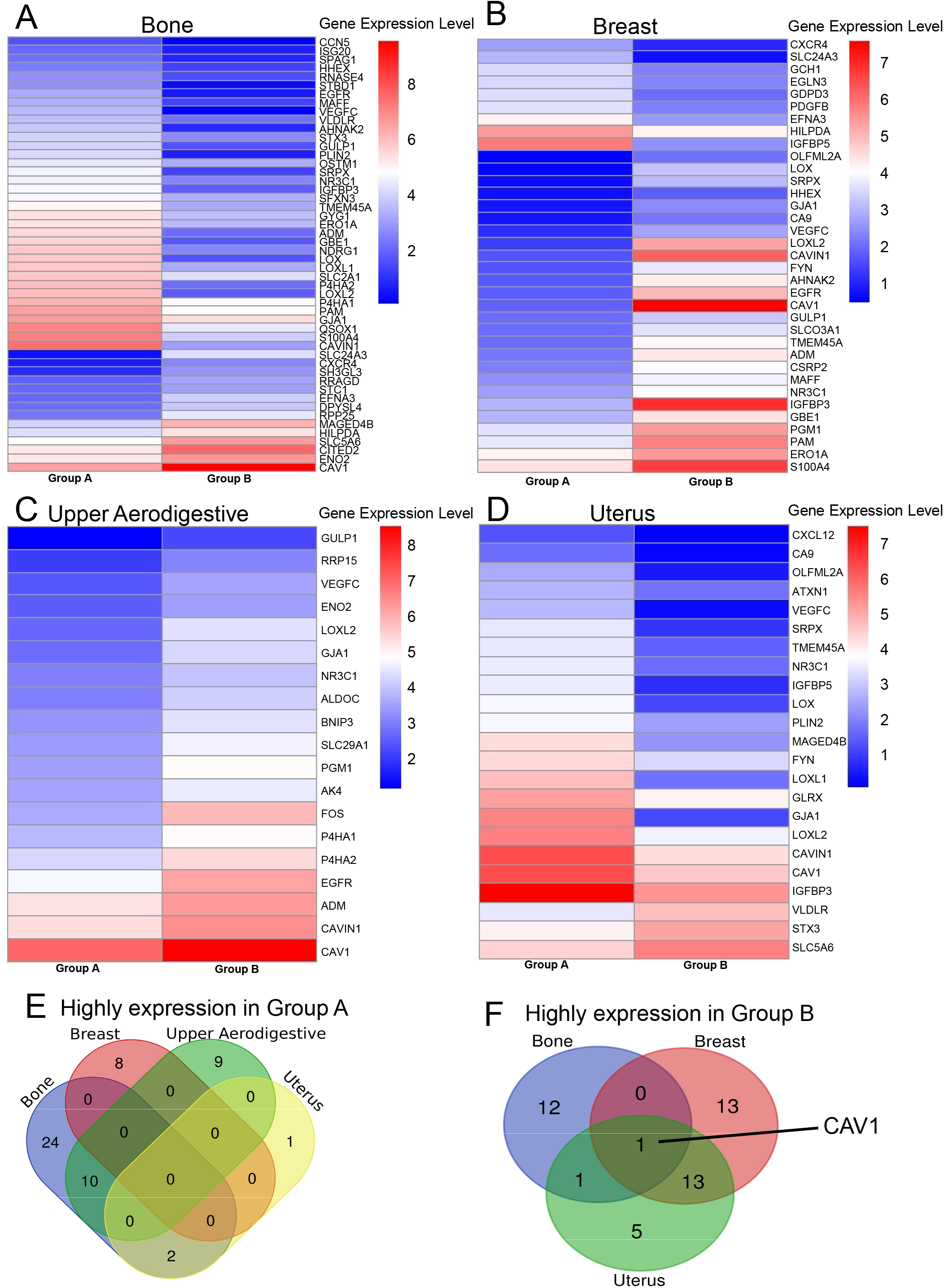
Extraction of DEGs in reclassified cancer cell lines. DEGs in Groups A and B were extracted following |logFC|>1. The average expression levels for each group are shown in the heatmap. The number of genes upregulated in Groups A and B are summarized in the Venn diagram.

## Discussion

HIF1 is a transcription factor that regulates the transcription of genes involved in tumor malignancy. This study examined the classification of cancer cell lines by HIF1 downstream gene expressions. Results identified that cell lines in 24 lineages were classified into two clusters, four of which revealed a difference in metastatic potential. Interestingly, DEGs are quite different among those tumor types, strongly suggesting a unique metastatic mechanism although HIF1 is commonly involved in the metastasis. Additionally, such tumor-specific HIF1 downstream gene set may provide a new tool for classification with different metastatic potentials and prognoses.

This study used HIF1 downstream gene, including genes that have been investigated concerning metastasis. CAV1 [20–22] is highly expressed in breast, bone, and upper aerodigestive cancers in Group B, and uterine cancer in Group A. C-X-C Motif Chemokine Receptor 4 (CXCR4) [23–26] is highly expressed in bone tumors in Group B and breast cancer in Group A, with no expression variation in the other two cancers. This is inconsistent with the study reporting that CXCR4 acts in metastasis-promoting breast cancer. Our study was inconsistent with previous reports on the association between differential CAV1, CXCR4, and metastatic potential expression by these genes. Additional genes should be analyzed to understand the metastatic mechanism of each tumor type.

This study classified subgroups with different metastatic potentials using HIF1 downstream genes. Our study has the following limitations. First, only 486 cell lines were available for metastatic potential analysis. Therefore, evaluating the statistical significance in most cases and excluding the bias of cancer cell lines was difficult. Next, we have not confirmed the activity of HIF1 in cancer cell lines. Finally, not only HIF1 but also HIF2 plays an important role as a transcription factor in hypoxia [27]. Some genes are activated by HIF1 and HIF2, but others are regulated differently by the two, although both act as transcription factors [27]. Marker gene identification from HIF1 downstream genes in each class of tumor contributes to the diagnosis of metastatic potential and treatment despite such study limitations.

## Materials and methods

### Data collection

The HIF1 downstream gene list was downloaded from MSigDB ([19]) in the form of “ELVIDGE_HIF1A_TARGETS_DN,” “ELVIDGE_HIF1A_TARGETS_UP” [18].

The RNA-seq dataset [16] and the metastatic potential score dataset [17] of the cancer cell lines were downloaded from the Depmap portal (https://depmap.org/portal/) in December 2022. Gene expressions are provided after log2 transformation, using a pseudo-count of 1; log2 (Transcripts per million [TPM]+1). The dataset of cell lines was analyzed with annotation data.

### Clustering and extract DEGs

Hierarchical clustering was used to identify distinct subgroups among cancer cell lines using the HIF1 downstream gene expression. The hierarchical clustering was performed based on the Euclidean distance following Ward’s method. The threshold value for DEGs was set at the absolute value of log two FC of >1.0.

### Statistical analysis

A Mann–Whitney U test (two-tailed) was used for significant analyses.

P-values of□<0.05 was considered statistically significant. Mann–Whitney U test was performed in R (version 4.1.3) [28].

### Data visualization

The heatmap and violin plot was drawn by the “pheatmap” “vioplot” package using the R statistical programming language[29, 30].

## Data availability

The results shown here are based upon data generated by CCLE MSigDB and are available in a public repository from the https://depmap.org/portal/, http://www.gsea-msigdb.org/gsea/index.jsp websites.

## Acknowledgments

We are grateful to all members of the lab for stimulating discussions during the manuscript preparation.

## Author contributions

All authors have participated in the design of the study. KN performed the bioinformatical analyses. KN and JN wrote the manuscript. All the authors reviewed and edited the manuscript.

## Figure legends

Table 1. Specific downstream genes in each cancer type

**Supplementary Figure S1 Subtype of cancer cell lines in CCLE**

The subtype information for each of the cell lines added by CCLE was shown by color labels.

**Supplementary Figure S2. Expression of DEGs in reclassified cancer cell lines**.

The expression of DEGs was aligned based on the hierarchical clustering results of the cell lines in Figure 1, and the expression of only DEGs was shown in the heatmap.

**Supplementary Table S1**. Classified cancer cell line list in the clustering of HIF1 downstream genes

**Supplementary Table S2**. Classified cancer cell lines in the clustering with the metastatic score

**Supplementary Table S3**. DEGs of reclassified cancer cell lines

## Notes

### Competing Interest Statement

The authors have declared no competing interest.

