## Supplementary material for "Metastatic potentials classified with hypoxia-inducible factor 1 downstream gene in pan-cancer cell lines": Table

| Table.1 Specific downstream genes in each cancer type | | | | | | | | | | | | |
| --- | --- | --- | --- | --- | --- | --- | --- | --- | --- | --- | --- | --- |
| High expression genes in Group A | | | | | | | High expression genes in Group B | | | | | |
| Bone and Upper aerodigestive | Bone and Uterus | Bone | | Breast | Upper aerodigestive | Uterus | Bone and Breast and Uterus | Bone and Uterus | Breast and Uterus | Bone | Breast | Uterus |
| EGFR | STX3 | TMEM45A | MAFF | GDPD3 | AK4 | SLC5A6 | CAV1 | MAGED4B | TMEM45A | STC1 | CSRP2 | GLRX |
| P4HA1 | VLDLR | STBD1 | GBE1 | GCH1 | RRP15 |  |  |  | OLFML2A | RRAGD | AHNAK2 | PLIN2 |
| CAVIN1 |  | OSTM1 | GYG1 | HILPDA | FOS |  |  |  | LOX | RPP25 | EGFR | LOXL1 |
| ADM |  | LOX | SRPX | CXCR4 | SLC29A1 |  |  |  | IGFBP3 | DPYSL4 | S100A4 | ATXN1 |
| LOXL2 |  | AHNAK2 | PAM | PDGFB | BNIP3 |  |  |  | CAVIN1 | SLC5A6 | ERO1A | CXCL12 |
| VEGFC |  | SLC2A1 | LOXL1 | EFNA3 | CAV1 |  |  |  | FYN | CXCR4 | MAFF |  |
| GULP1 |  | IGFBP3 | SFXN3 | EGLN3 | PGM1 |  |  |  | LOXL2 | HILPDA | GBE1 |  |
| NR3C1 |  | ISG20 | SPAG1 | SLC24A3 | ALDOC |  |  |  | SRPX | SH3GL3 | ADM |  |
| GJA1 |  | QSOX1 | RNASE4 |  | ENO2 |  |  |  | VEGFC | EFNA3 | PAM |  |
| P4HA2 |  | S100A4 | HHEX |  |  |  |  |  | CA9 | CITED2 | PGM1 |  |
|  |  | ERO1A | CCN5 |  |  |  |  |  | NR3C1 | ENO2 | GULP1 |  |
|  |  | NDRG1 |  |  |  |  |  |  | IGFBP5 | SLC24A3 | HHEX |  |
|  |  | PLIN2 |  |  |  |  |  |  | GJA1 |  | SLCO3A1 |  |
