## Supplementary figures and images for "Metastatic potentials classified with hypoxia-inducible factor 1 downstream gene in pan-cancer cell lines"

### Supplementary Figures

Supplementary Figure S1

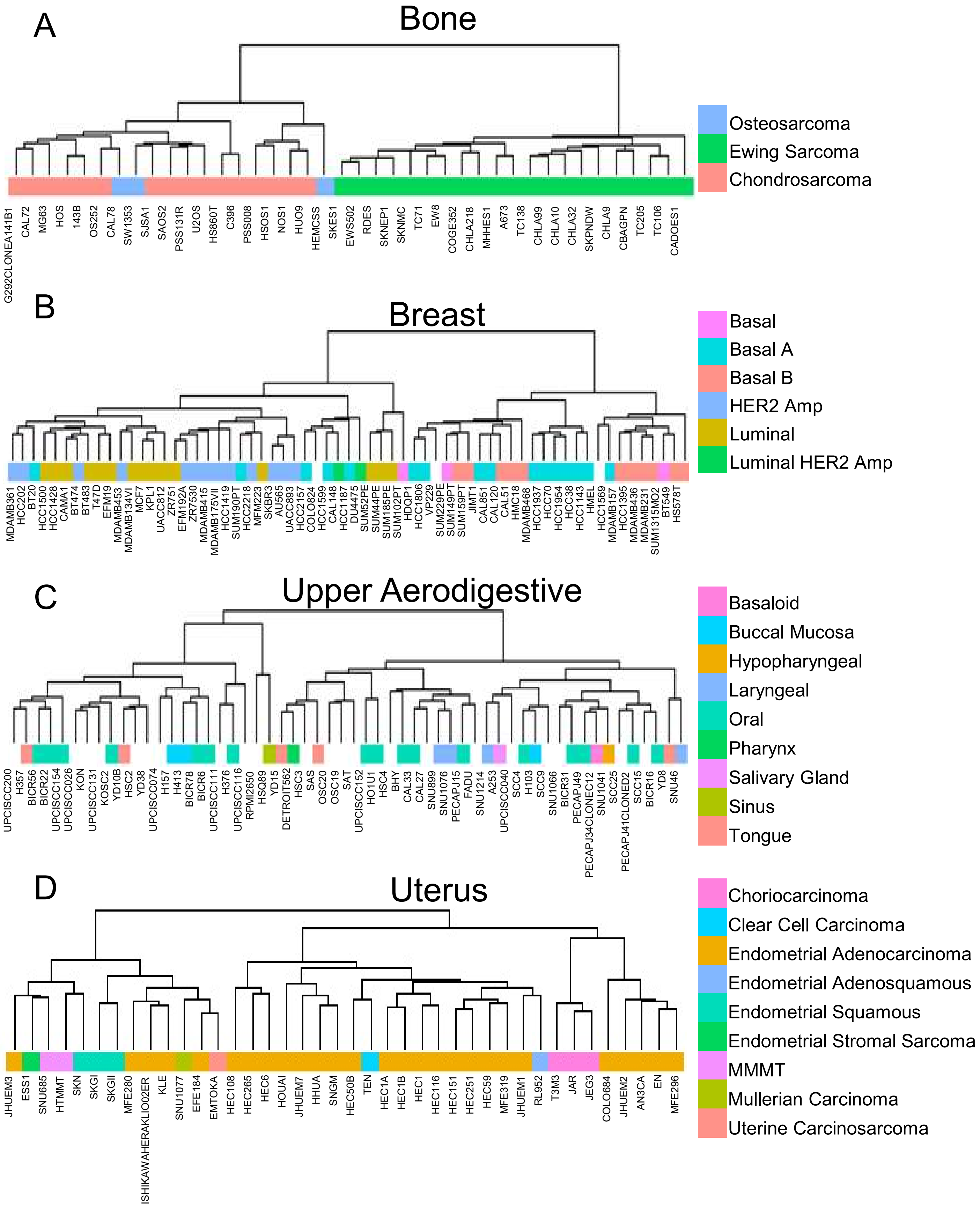

Supplementary FigureS2

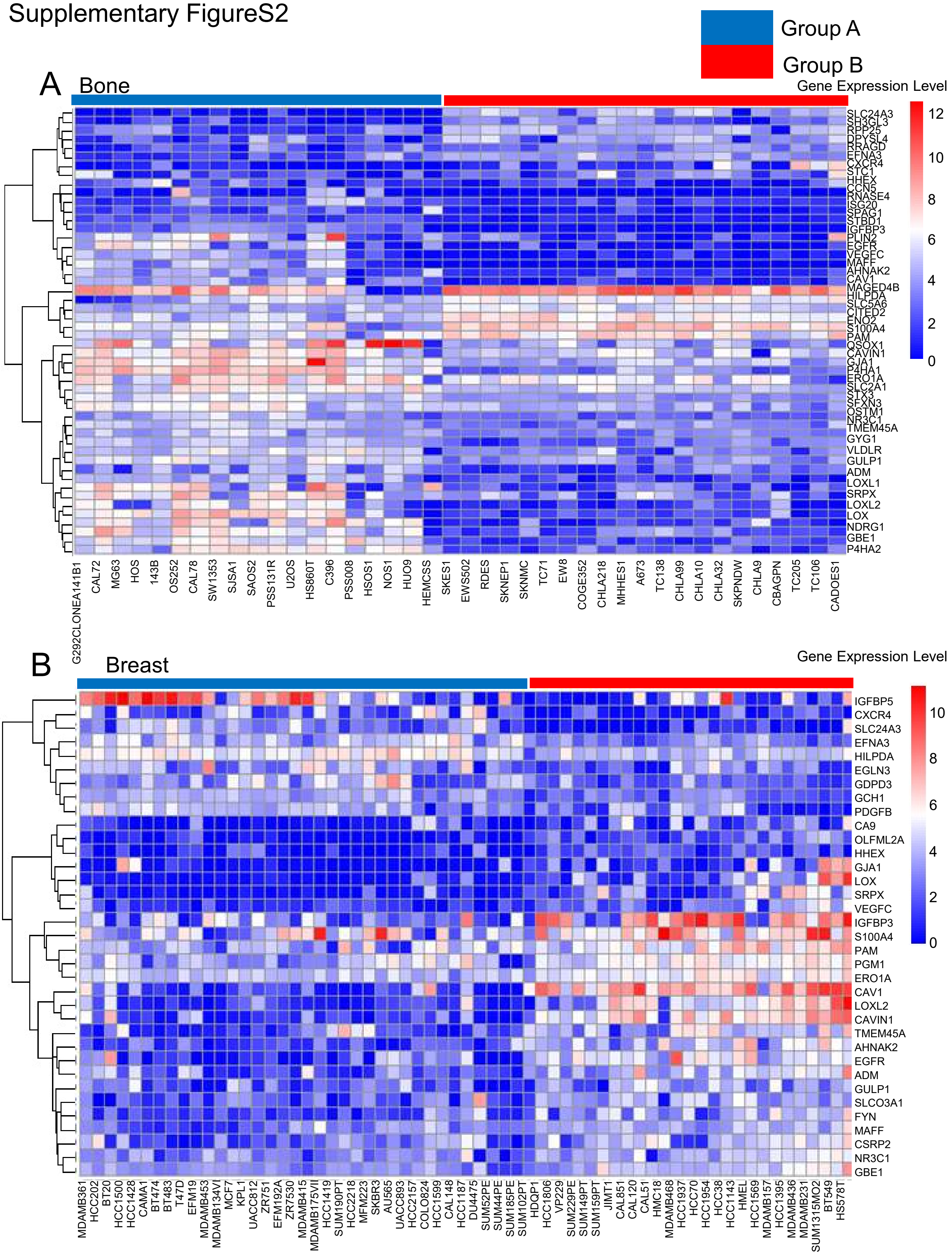

Supplementary FigureS2

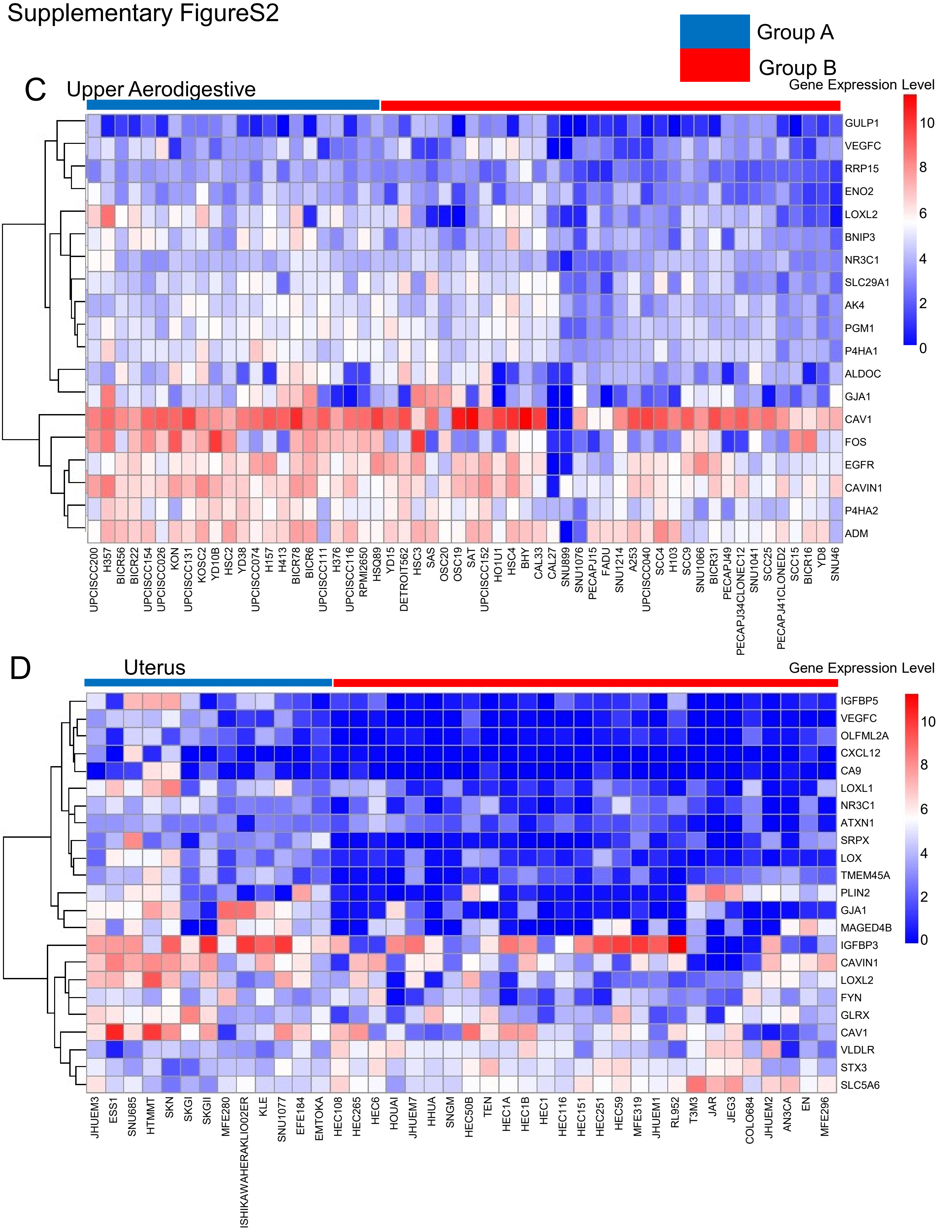
