## Supplementary Tables for "Metastatic potentials classified with hypoxia-inducible factor 1 downstream gene in pan-cancer cell lines"

| Supplementary Table S1. Classified cancer cell line list in the clustering of HIF1 downstream genes | | | | | | | |
| --- | --- | --- | --- | --- | --- | --- | --- |
| Bone | | Breast | | Upper aerodigestive | | Uterus | |
| Group A | Group B | Group A | Group B | Group A | Group B | Group A | Group B |
| 143B | A673 | AU565 | BT549 | BICR22 | A253 | EFE184 | AN3CA |
| C396 | CADOES1 | BT20 | CAL120 | BICR56 | BHY | EMTOKA | COLO684 |
| CAL72 | CBAGPN | BT474 | CAL51 | BICR6 | BICR16 | ESS1 | EN |
| CAL78 | CHLA10 | BT483 | CAL851 | BICR78 | BICR31 | HTMMT | HEC1 |
| G292CLONEA141B1 | CHLA218 | CAL148 | HCC1143 | H157 | CAL27 | ISHIKAWAHERAKLIO02ER | HEC108 |
| HEMCSS | CHLA32 | CAMA1 | HCC1395 | H357 | CAL33 | JHUEM3 | HEC116 |
| HOS | CHLA9 | COLO824 | HCC1569 | H376 | DETROIT562 | KLE | HEC151 |
| HS860T | CHLA99 | DU4475 | HCC1806 | H413 | FADU | MFE280 | HEC1A |
| HSOS1 | COGE352 | EFM19 | HCC1937 | HSC2 | H103 | SKGI | HEC1B |
| HUO9 | EW8 | EFM192A | HCC1954 | HSQ89 | HO1U1 | SKGII | HEC251 |
| MG63 | EWS502 | HCC1187 | HCC38 | KON | HSC3 | SKN | HEC265 |
| NOS1 | MHHES1 | HCC1419 | HCC70 | KOSC2 | HSC4 | SNU1077 | HEC50B |
| OS252 | RDES | HCC1428 | HDQP1 | RPMI2650 | OSC19 | SNU685 | HEC59 |
| PSS008 | SKES1 | HCC1500 | HMC18 | UPCISCC026 | OSC20 |  | HEC6 |
| PSS131R | SKNEP1 | HCC1599 | HMEL | UPCISCC074 | PECAPJ15 |  | HHUA |
| SAOS2 | SKNMC | HCC202 | HS578T | UPCISCC111 | PECAPJ34CLONEC12 |  | HOUAI |
| SJSA1 | SKPNDW | HCC2157 | JIMT1 | UPCISCC116 | PECAPJ41CLONED2 |  | JAR |
| SW1353 | TC106 | HCC2218 | MDAMB157 | UPCISCC131 | PECAPJ49 |  | JEG3 |
| U2OS | TC138 | KPL1 | MDAMB231 | UPCISCC154 | SAS |  | JHUEM1 |
|  | TC205 | MCF7 | MDAMB436 | UPCISCC200 | SAT |  | JHUEM2 |
|  | TC71 | MDAMB134VI | MDAMB468 | YD10B | SCC15 |  | JHUEM7 |
|  |  | MDAMB175VII | SUM1315MO2 | YD38 | SCC25 |  | MFE296 |
|  |  | MDAMB361 | SUM149PT |  | SCC4 |  | MFE319 |
|  |  | MDAMB415 | SUM159PT |  | SCC9 |  | RL952 |
|  |  | MDAMB453 | SUM229PE |  | SNU1041 |  | SNGM |
|  |  | MFM223 | VP229 |  | SNU1066 |  | T3M3 |
|  |  | SKBR3 |  |  | SNU1076 |  | TEN |
|  |  | SUM102PT |  |  | SNU1214 |  |  |
|  |  | SUM185PE |  |  | SNU46 |  |  |
|  |  | SUM190PT |  |  | SNU899 |  |  |
|  |  | SUM44PE |  |  | UPCISCC040 |  |  |
|  |  | SUM52PE |  |  | UPCISCC152 |  |  |
|  |  | T47D |  |  | YD15 |  |  |
|  |  | UACC812 |  |  | YD8 |  |  |
|  |  | UACC893 |  |  |  |  |  |
|  |  | ZR751 |  |  |  |  |  |
|  |  | ZR7530 |  |  |  |  |  |

| Supplementary Table S2. Classified cancer cell lines in the clustering with metastatic score | | | | | | | |
| --- | --- | --- | --- | --- | --- | --- | --- |
| Bone | | Breast | | Upper aerodigestive | | Uterus | |
| Group A | Group B | Group A | Group B | Group A | Group B | Group A | Group B |
| CAL78 | A673 | BT474 | BT549 | BICR56 | BICR16 | ESS1 | AN3CA |
| G292CLONEA141B1 | CADOES1 | CAMA1 | CAL120 | BICR6 | BICR31 | ISHIKAWAHERAKLIO02ER | EN |
| HOS | EWS502 | EFM192A | CAL51 | HSC2 | CAL27 | MFE280 | HEC108 |
| MG63 | MHHES1 | HCC1419 | HCC1143 | YD10B | DETROIT562 | SNU1077 | HEC151 |
| SJSA1 | SKES1 | HCC1428 | HCC1806 | YD38 | FADU | SNU685 | HEC1A |
| SW1353 | SKNEP1 | KPL1 | HCC1937 |  | HSC3 |  | HEC1B |
| U2OS |  | MCF7 | HCC38 |  | PECAPJ15 |  | HEC251 |
|  |  | MDAMB175VII | HDQP1 |  | PECAPJ41CLONED2 |  | HEC265 |
|  |  | T47D | HMC18 |  | PECAPJ49 |  | HEC59 |
|  |  | ZR751 | JIMT1 |  | SCC25 |  | HEC6 |
|  |  |  | MDAMB231 |  | SNU1041 |  | JHUEM2 |
|  |  |  | MDAMB436 |  | SNU1066 |  | MFE296 |
|  |  |  | MDAMB468 |  | SNU1076 |  | MFE319 |
|  |  |  |  |  | SNU1214 |  | RL952 |
|  |  |  |  |  | SNU46 |  | SNGM |
|  |  |  |  |  | YD15 |  | TEN |
|  |  |  |  |  | YD8 |  |  |

| Supplementary Table S3. DEGs of reclassified cancer cell lines | | | | | |
| --- | --- | --- | --- | --- | --- |
| Bone | | Breast | | Uterus | Upper aerodigestive |
| CCN5 | LOX | CXCR4 | ADM | VLDLR | GULP1 |
| ISG20 | LOXL1 | SLC24A3 | CSRP2 | STX3 | RRP15 |
| SPAG1 | SLC2A1 | GCH1 | MAFF | SLC5A6 | ENO2 |
| HHEX | P4HA2 | EGLN3 | NR3C1 | CXCL12 | VEGFC |
| RNASE4 | LOXL2 | GDPD3 | IGFBP3 | CA9 | NR3C1 |
| STBD1 | P4HA1 | PDGFB | GBE1 | VEGFC | ALDOC |
| EGFR | PAM | EFNA3 | PGM1 | OLFML2A | GJA1 |
| MAFF | GJA1 | HILPDA | PAM | IGFBP5 | BNIP3 |
| VEGFC | QSOX1 | IGFBP5 | ERO1A | SRPX | LOXL2 |
| VLDLR | S100A4 | OLFML2A | S100A4 | LOX | AK4 |
| AHNAK2 | CAVIN1 | LOX |  | GJA1 | SLC29A1 |
| STX3 | SLC24A3 | SRPX |  | TMEM45A | PGM1 |
| GULP1 | CXCR4 | HHEX |  | NR3C1 | P4HA1 |
| PLIN2 | SH3GL3 | GJA1 |  | ATXN1 | P4HA2 |
| OSTM1 | RRAGD | CA9 |  | LOXL1 | FOS |
| SRPX | STC1 | VEGFC |  | MAGED4B | EGFR |
| NR3C1 | EFNA3 | LOXL2 |  | PLIN2 | ADM |
| IGFBP3 | DPYSL4 | CAVIN1 |  | FYN | CAVIN1 |
| SFXN3 | RPP25 | FYN |  | LOXL2 | CAV1 |
| TMEM45A | MAGED4B | AHNAK2 |  | GLRX |  |
| GYG1 | HILPDA | EGFR |  | CAVIN1 |  |
| ERO1A | SLC5A6 | CAV1 |  | CAV1 |  |
| ADM | CITED2 | GULP1 |  | IGFBP3 |  |
| GBE1 | ENO2 | SLCO3A1 |  |  |  |
| NDRG1 | CAV1 | TMEM45A |  |  |  |
